# Integration of the hammerhead ribozyme into structured RNAs to measure ligand-binding events for riboswitch candidates and aptamers

**DOI:** 10.1101/2025.04.10.648167

**Authors:** Shenglan Zhang, Yinghong Lin, Ting Gao, Binfen Chen, Weibin Wu, Shanshan Fang, Kexin Fan, Yuqing Lai, Yezi Lin, Rongqin Ke, Sanshu Li

## Abstract

Some structured RNAs, such as riboswitches and aptamers, can bind their cognate ligands and have been used in biosensors and gene expression control elements. However, current methods for detecting ligand binding to structured RNAs are either severely limited or inconvenient. In this study, we design a multi-base pair bridge to integrate a hammerhead ribozyme into structured RNAs to detect ligand binding events. The experimental results demonstrate that the length of the bridge has a significant effect on the cleavage of the ribozyme; optimal cleavage can be achieved with three to six base pairs in the bridge. The dissociation constant (*K*_D_) values obtained through this method are in agreement with those determined by in-line probing techniques, and 1 pmol of allosteric ribozyme RNA is sufficient for measurement. We apply this method to evaluate the binding affinity of the riboswitch candidate Motif_9307. Our findings indicate that this motif has no binding affinity for S-adenosylmethionine or several other tested ligands, which is consistent with the results of the in-line probing experiments. Notably, our method reveals an increase in cleavage activity when yeast extract is added as a mixture of ligands, suggesting that the ligand of Motif_9307 is present in the extract. In conclusion, we develop an alternative approach for measuring ligand binding events associated with riboswitch candidates and aptamers.

## Introduction

Riboswitches are structured non-coding RNAs that are typically located in the 5’ untranslated regions (UTRs) of mRNAs, where they fold and bind to metabolites or other small molecules to regulate gene expression[1, 2]. Riboswitches are actually aptamers carrying expression platforms. Aptamers are single-stranded DNA or RNA molecules that can fold into structures and bind to ligands with high specificity and affinity[3-7]. They have been engineered as various sensitive biosensors and nucleic acid drugs[8-14].

The aptamer and expression platform of riboswitches are two overlapping components[2, 15]. The aptamer can bind to ligands, and the expression platform can turn on or off gene expression. Examples of expression platforms include ribosomal binding sites (RBSs), rho-independent terminators, and splicing site proximal regions[1, 2, 16-18]. The conformation of the aptamer changes upon ligand binding, resulting in exposure or sequestration of the expression platform. In addition, tandem riboswitches, such as the guanine aptamer combined with the phosphoribosyl pyrophosphate (PRPP) aptamer, can function as an IMPLY Boolean logic gate to regulate the transcription of messenger RNAs for purine biosynthesis in bacteria[19], implying their possible applications as logic control circuits[19, 20]. Hence, these structured RNAs serve as genetic switches that can activate or repress gene expression in response to changes in the levels of their target ligands[1, 17, 21].

Riboswitches are widely distributed in bacteria and archaea[2, 18, 22-24]. The thiamine pyrophosphate (TPP) riboswitch also appears in eukaryotes, such as algae, plants, and fungi[25-30]. In general, riboswitches participate in the regulation of all types of genes, including metabolite genes, ion transporter genes, cofactor synthesis genes, virulence-related genes, and signaling pathway-related genes[15, 22, 31]. Thus far, approximately 56 classes of riboswitches have been identified[1, 22]. Riboswitch classes recognize a large variety of ligands, including ribonucleotide derivatives (e.g., xanthine, preQ_1_, and 2’-dG), ribonucleotide precursors (e.g., guanine, adenine, 5-aminoimidazole-4-carboxamide riboside 5’-triphosphate (ZTP), and ADP), cofactors (TPP, adenosylcobalamin (adoCbl), S-adenosylmethionine (SAM), and nicotinamide adenine dinucleotide (NAD^+^)), amino acids (e.g., glycine, lysine, and glutamine), elemental ions (Mn^2+^, Fluoride, Mg^2+^, Li^+^, and Na^+^), signaling molecules (e.g., c-di-GMP, c-di-AMP, and ppGpp), glucose-related molecules (glucosamine-6-phosphate), and others (e.g., guanidine and azaaromatic compounds) [1, 22]. The TPP and AdoCbl riboswitches are the most common, with 16,701 and 14,646 representatives, respectively[1].

Among these riboswitches, the glmS riboswitch is special in that it is also a ribozyme, a naturally occurring self-cleaving RNA motif that catalyzes site-specific RNA cleavage reactions[32]. Thus far, more than 13 classes of natural ribozymes have been discovered, including group I and group II introns[33], RNase P, Ribosomal RNA, Hammerhead, Hairpin, hepatitis delta virus (HDV), Neurospora VS, glmS, Twister, Twister sister, Hatchet, and Pistol. The binding of glucosamine-6-phosphate to the glmS riboswitch causes the embedded ribozyme to self-cleave[34]. The c-di-GMP-II riboswitch uses the group I intron (a ribozyme) as its expression platform to regulate gene expression[35, 36]. It seems that, in nature, riboswitches preferentially use ribozymes as their expression platforms, alongside ribosome binding sites and Rho-independent terminators.

Drawing on our understanding of natural riboswitches and ribozymes, researchers selected artificial riboswitches to sense ligands by Systematic Evolution of Ligands by Exponential Enrichment (SELEX). For example, Dr. Breaker’s group selected allosteric hammerhead ribozymes that can sense certain divalent metal ions[37], theophylline[38], and c-di-GMP[39] from random sequences. Similarly, Gu’s laboratory also selected artificial riboswitches to fabricate a biosensor for detecting TPP concentrations in blood [40]. Moreover, Hartig’s group employed site-directed mutagenesis to create an allosteric ribozyme to regulate the translation of mammalian mRNAs[41]. Recently, we integrated DNAzymes (single stranded DNA molecules that can self-cleave) into DNA aptamers in the SELEX cycles to select allosteric DNAzymes, which can sense different ligands and induce DNA self-cleavage[42, 43]. Previously, we applied bioinformatics analysis to search metagenomes and discovered thousands of novel structured noncoding RNAs, including hundreds of riboswitch candidates. However, we encountered difficulties finding a convenient and sensitive in vitro approach to validate these riboswitch candidates. Currently, several in vitro approaches are used to validate riboswitches; these include in-line probing[44], isothermal titration calibration (ITC)[45, 46], and the SYBR Green fluorometric assay[47]. In-line probing is the most powerful in vitro method for testing whether structured noncoding RNA can bind to a ligand and whether the binding is specific and meaningful in a genomic context. However, this approach requires the use of a radionuclide (^32^P) to be sufficiently sensitive to observe hundreds of labeled RNA bands on long polyacrylamide gels, and the use of radionuclides is not convenient for commonly equipped laboratories. In addition, radionuclides are expensive, short-lived, and potentially hazardous to users. ITC can be used to detect binding events based on the change in heat upon ligand binding; however, this method requires a large amount of RNA (approximately 100 μl of 0.2 mM RNA) and a large number of ligands to allow the generation of a measurable temperature shift. The SYBR Green fluorometric assay can also be used for ligand-binding analysis based on the competition between SYBR Green and the ligand; however, this method is less sensitive than in-line probing and ITC[46].

In this study, we designed a bridge to integrate the hammerhead ribozyme into structured RNAs to detect ligand-binding events. The bridge is composed of a few base pairs derived from stem II of the hammerhead ribozyme and a few base pairs from stem P1 of aptamers or riboswitches. We tested this approach by fusing a theophylline aptamer and two TPP riboswitches with the hammerhead ribozyme through bridges with a few base pairs to observe and measure the cleavage of allosteric ribozymes. Based on the results of the binding specificity and affinity (the *K*_D_ values) of these allosteric ribozymes, we found that this method was both sensitive and convenient. We also used this method to analyze a previously identified riboswitch candidate using our bioinformatics approach. The findings demonstrated its ability to bind to the ligand and induce cleavage of the integrated ribozyme. When combined with in vivo data, we propose that the riboswitch candidate Motif_9307 functions as a regulatory RNA that senses specific metabolites within the cell, excluding SAM itself. Consequently, this alternative approach could be used to assess ligand-binding events for riboswitch candidates and aptamers.

## Materials and Methods

### Preparation of DNA template for allosteric ribozymes

A DNA template containing the T7 promoter, aptamers (or riboswitches) and hammerhead ribozyme was prepared by PCR. In the PCR, two overlapping long primers (Supplementary Table S1) were used to generate a long DNA product, using a method called overlapping PCR to prepare full-length DNA as described in the paper[48]. Specifically, the overlapping primers ThiC-TPP I-forward and ThiC-TPP I-reverse (overlapping region 5′ GAAATACCCGTATCACCTGATCTGG) were used to generate the DNA template for the ThiC-TPP allosteric ribozyme. First, prepare the PCR reaction containing 25 μL of 2× Accurate Taq Master Mix (from Accurate Biology, Hunan, China), 2 μL each of forward and reverse primers (10 μM) and 21 μL of H_2_O. The reaction is then performed in a PCR amplifier according to the following cycling conditions 94°C for 30 s, followed by 30 cycles of 98°C for 10 s, 52°C for 30 s, 72°C for 15 s, with a final extension at 72°C for 2 min. After the PCR cycles, the full-length DNA products were purified using a PCR purification kit (Tiangen Biotech, Shanghai, China). DNA templates for other allosteric ribozymes were also prepared using the same procedure.

### Preparation of RNA for allosteric ribozymes

We transcribed the full-length DNA template into RNA using a transcription kit (Vazyme Biotech, Nanjing, China). The full-length RNA was separated by 10% urea-denaturing polyacrylamide gel electrophoresis (PAGE), and the gel pieces were crushed, soaked in elution buffer (Tris-HCl 10 mM, pH 7.5 at 23°C, 200 mM NaCl, and 1 mM EDTA), and recovered by ethanol precipitation.

### Allosteric ribozyme cleavage assays

Cleavage assays were performed as previously described[49, 50]. Briefly, approximately 1 pmol of RNA was combined with cleavage buffer (30 mM HEPES [pH 7.5], 100 mM KCl) in a volume of 17.6 μL. The mixture was heated at 80°C for 1 min and then allowed to cool to room temperature for 10 min. Subsequently, 0.4 μL of 1 M MgCl_2_ (resulting in a final concentration of 20 mM) and 2 μL of the ligand (to achieve the required concentration) were added and mixed thoroughly using a pipette. The tubes were then incubated at 25°C for 30 min. After the reaction, the tubes were added with 20 μL of 2X loading buffer (10 M Urea, 1.5 mM EDTA [pH 8.0], 0.05% xylene cyanole, and 0.05% bromophenol blue) and mixed well with a pipette. Cleavage products were separated on 10% PAGE gels and visualized by staining with SYBR Gold (Thermo Fisher Scientific, Eugene, USA). The cleavage fraction was determined as the intensity of the 5’ cleavage RNA products divided by the total RNA.

### Dissociation constant (*K*_D_) measurements

The apparent *K*_D_ values were determined using a previously described method[42, 43, 48]. The detailed steps for the *K*_D_ assay are consistent with those described above for allosteric ribozyme cleavage assays, in which varying concentrations of ligands were added to the reactions. The experiments were repeated three times. The band intensity was quantified using ImageJ[51], with a consistently defined area applied to all bands. The cleavage ratio was calculated as the intensity of the 5′ cleavage band divided by the total intensity of both the 5′ cleavage band and the full-length band. The cleavage ratios were normalized by dividing each ratio by the maximum cleavage ratio obtained, and the resulting value was subsequently multiplied by 100. The concentration of the ligand that induces a 50% shift in the normalized cleavage activity of the allosteric ribozyme is referred to as the apparent *K*_D_. The *K*_D_ value was calculated using GraphPad Prism with the function of specific binding with Hill slope 1.0 and the equation Y = B_max_ × X/(*K*_D_ + X), where B_max_ is the maximum-specific binding.

### In-line probing

The assay was conducted following previously established protocols[27, 48]. In brief, 5’ ^32^P-labeled RNAs were incubated with various concentrations of ligands at 25°C for 36 h in the presence of 100 mM KCl, 50 mM Tris-HCl (pH 8.3 at 23°C), and 20 mM MgCl_2_. The RNA spontaneous cleavage products were separated by denaturing (8 M urea) 10% polyacrylamide gel electrophoresis (PAGE). The images were captured using a PhosphorImager system (GE Healthcare, Chicago, USA).

### In vivo reporter construction

The plasmid pBS1C*lacZ*[52], obtained from Addgene, contains a *lacZ* reporter, *amyE* integration site, and resistance genes *amp*^*R*^ and *cm*^*R*^. The *lysC* promoter (5’GCAAAAATAATGTTGTCCTTTTAAATAAGATCTGATAAAATGTGAACTAAT) was inserted between the EcoRI and BamHI sites in the plasmid. Additionally, HindIII was incorporated in front of the *lacZ* reporter (Table S1). Subsequently, the DNA fragment encoding Motif_9307 RNA was inserted between BamHI and HindIII to create the final plasmid construct (pBS1C-motif_9307), which featured an in-frame translational fusion of Motif_9307 with the adjacent *lacZ* gene.

### Integration of the Motif_9307-lacZ reporter-fusion construct into *B. subtilis*

The plasmid pBS1C motif_9307 was transfected into the *B. subtilis* strain PY79, as previously described[48]. Successful transformants were identified by selecting colonies that were chloramphenicol-resistant and ampicillin-sensitive. PCR was also used for confirmation using two pairs of primers listed in Table S1.

### *LacZ* (β-galactosidase) activity assay

Reporter gene *lacZ* activity assays were conducted as described previously[48]. Arbitrary units of β-galactosidase activity were calculated as fluorescence intensity divided by the total cell density (OD_595_) and reaction time.

### Identification of candidate structured noncoding RNA (ncRNA) Motif_9307 from archaea genomes using bioinformatics analyses

The structured ncRNA discovery pipeline was established as described in our previous paper[53]. Briefly, intergenic regions based on genomic annotations were extracted using our Perl scripts, BLAST[54] was used to align similar sequences to form clusters, and known ncRNAs and mis-annotated coding regions were removed from clusters. The remaining clusters were subjected to analysis with CMfinder[55] to identify variations and compatible mutations in the aligned sequences and predict the secondary structures of the RNA candidates. Motif_9307 is a structured ncRNA candidate. According to gene annotations from NCBI, the downstream genes associated with RNA Motif_9307 include *NosD* and its homologous genes, which encode a subunit of nitrous oxide reductase in the *nosDFYL tatE* operon structure[56]. The operon was activated in response to anaerobiosis and nitrate denitrification. The NosD subunit undergoes conformational changes upon ATP hydrolysis. Consequently, NosDFY alters its interaction partner, facilitating the transfer of copper ions from the chaperone NosL to the enzyme N_2_O reductase[57].

## Results

### Design of an allosteric ribozyme by integrating a hammerhead ribozyme into riboswitches and aptamers through a bridge for detecting ligand-binding events

Aptamers or riboswitches are single-stranded DNA or RNA molecules that can bind to cognate ligands. Binding affinity and specificity are important features of these ligand-binding elements. In this study, we developed a simple method to detect ligand-binding events by integrating a ribozyme into an aptamer. We designed bridges with a few base pairings to merge the aptamer with the ribozyme (Figure 1) rather than selecting bridges from random sequences using SELEX, which is a labor-intensive process. We designed bridges to transfer the ligand-binding information from the aptamer to the ribozyme, resulting in either the induction or inhibition of the self-cleavage of the ribozyme. Based on the SELEX results from previous experiments [37-39], we found that bridges composed of three to six base pairs might be adequately effective for the intended purpose. Initially, we designed the bridge to maintain one to three base pairs of stem II of the hammerhead unchanged, as well as one to three base pairs of stem P1 from either the aptamer or riboswitches.

**Figure 1.**
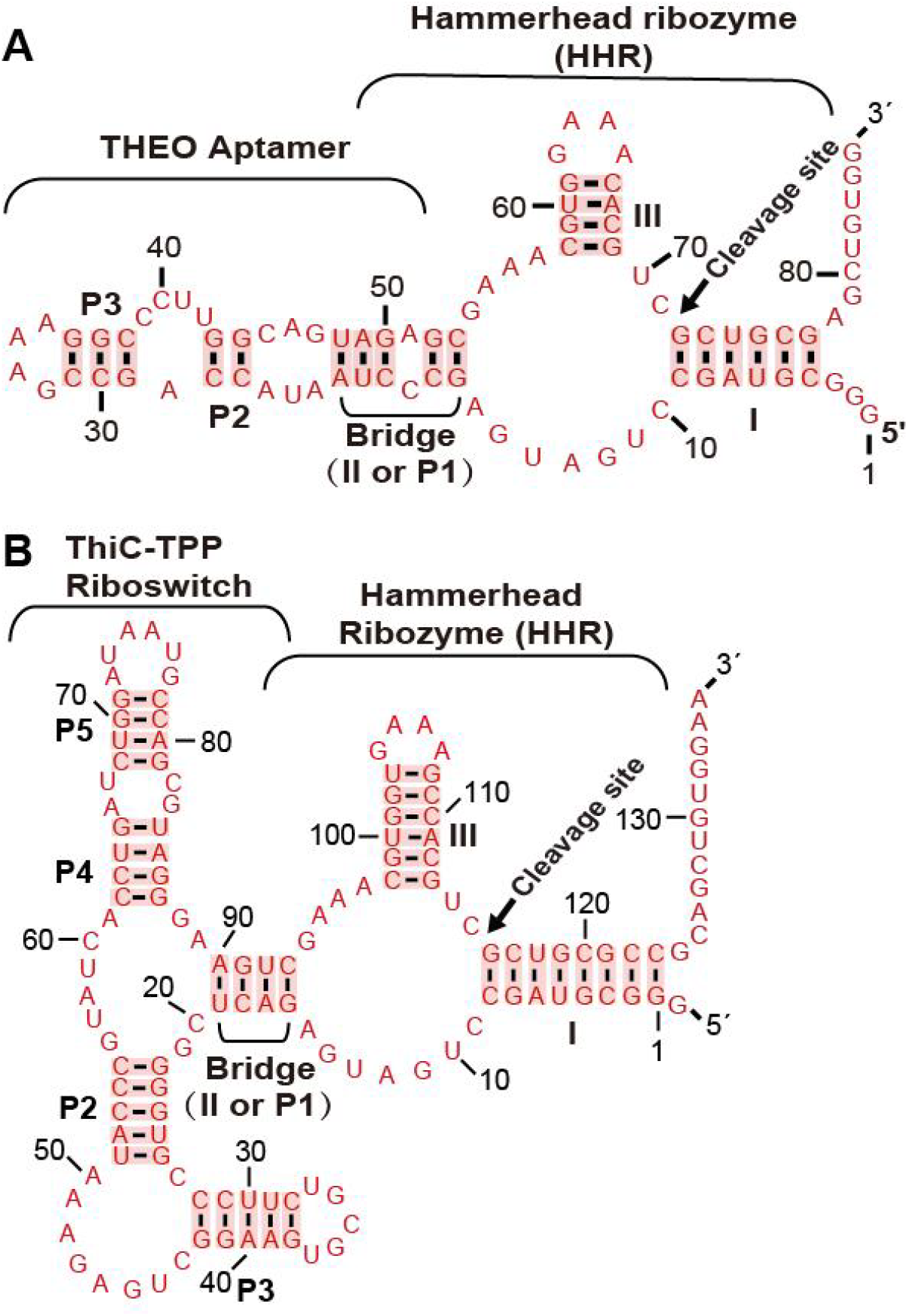
Combination of the hammerhead ribozymes with an aptamer or a riboswitch through a bridge. (A) The theophylline aptamer was merged with the hammerhead ribozyme through a bridge of five base pairs and an A–C bulge to separate the left part (three base pairs from stem P1 of the Theo aptamer) and the right part (two base pairs from stem II of the hammerhead ribozyme). (B) The TPP riboswitch from the bacterial *ThiC* gene was merged with the hammerhead ribozyme (named ThiC-TPP allosteric ribozyme). The bridge comprises four base pairs with one base pair from stem II of the ribozyme and three base pairs from stem P1 of the TPP riboswitch. The stems of the hammerhead ribozyme are designated as I–III. In contrast, the stems within the aptamer and riboswitches are denoted by P1–P5. The bridge was constructed from the partial sequences of stems P1 and II.

To demonstrate the working mechanism, we merged the theophylline aptamer[58] and TPP riboswitches from the bacterial *ThiC* gene[59] and from the fungal *NCU01977* gene[27] with the hammerhead ribozyme through bridges, resulting in three constructs called THEO allosteric ribozyme (Figure 1A), ThiC-TPP allosteric ribozyme (Figure 1B), and NCU-TPP allosteric ribozyme (Figure S1). Commonly, the hammerhead ribozyme performs self-cleavage as long as it can fold and form a proper tertiary structure. However, after the fusion of an aptamer with a ribozyme, the binding of the ligand changes the conformation of the aptamer. This conformation change twists the bridge to either facilitate or disrupt the ribozyme, affecting the self-cleavage of the ribozyme. Based on the changes in ribozyme self-cleavage, we can monitor the ligand-binding events of riboswitches and aptamers.

### Binding specificity of the allosteric ribozymes

To investigate the binding specificity of the THEO allosteric ribozyme, three closely related analogs were used for comparison (Figure 2A). Ribozyme cleavage assays showed that the binding of ligands to the THEO allosteric ribozyme promoted the cleavage of the ribozyme. Without a ligand, the THEO allosteric ribozyme produced a cleavage fraction of 0.163. Among these ligands, theophylline induced the most pronounced cleavage at a fraction of 0.451. The closely related analog 1-methylxanthine induced much less cleavage, with a fraction of 0.300, whereas 3-methylxanthine induced a cleavage fraction of 0.421, which was similar to that induced by theophylline (Figure 2B). These results indicate that the third position of the methyl group is important for ligand recognition because both theophylline and 3-methylxanthine contain this methyl group (Figure 2A). For the ThiC-TPP allosteric ribozyme with a bridge of four base pairs, several TPP analogs, including the closely related analog thiamine and the less closely related analog L-histidine, were used for comparison (Figure 2C). The addition of TPP inhibited the cleavage of the ThiC-TPP allosteric ribozyme. Without a ligand, the allosteric ribozyme itself produced a cleavage fraction of 0.403, whereas the addition of TPP reduced the cleavage fraction to 0.265. The TPP analog thiamine, which lacks a pyrophosphate group, reduced the cleavage fraction to 0.235, suggesting that both TPP and thiamine could be recognized by the allosteric ribozyme, probably through the pyrimidine ring and the thiazol group. However, the less closely related analog L-histidine, which contains an imidazole group, could barely inhibit self-cleavage, with a cleavage fraction of 0.393 (Figure 2D). These findings suggest that the ThiC-TPP allosteric ribozyme distinguishes between TPP and histidine and recognizes compounds that possess both a pyrimidine ring and a thiazole group.

**Figure 2.**
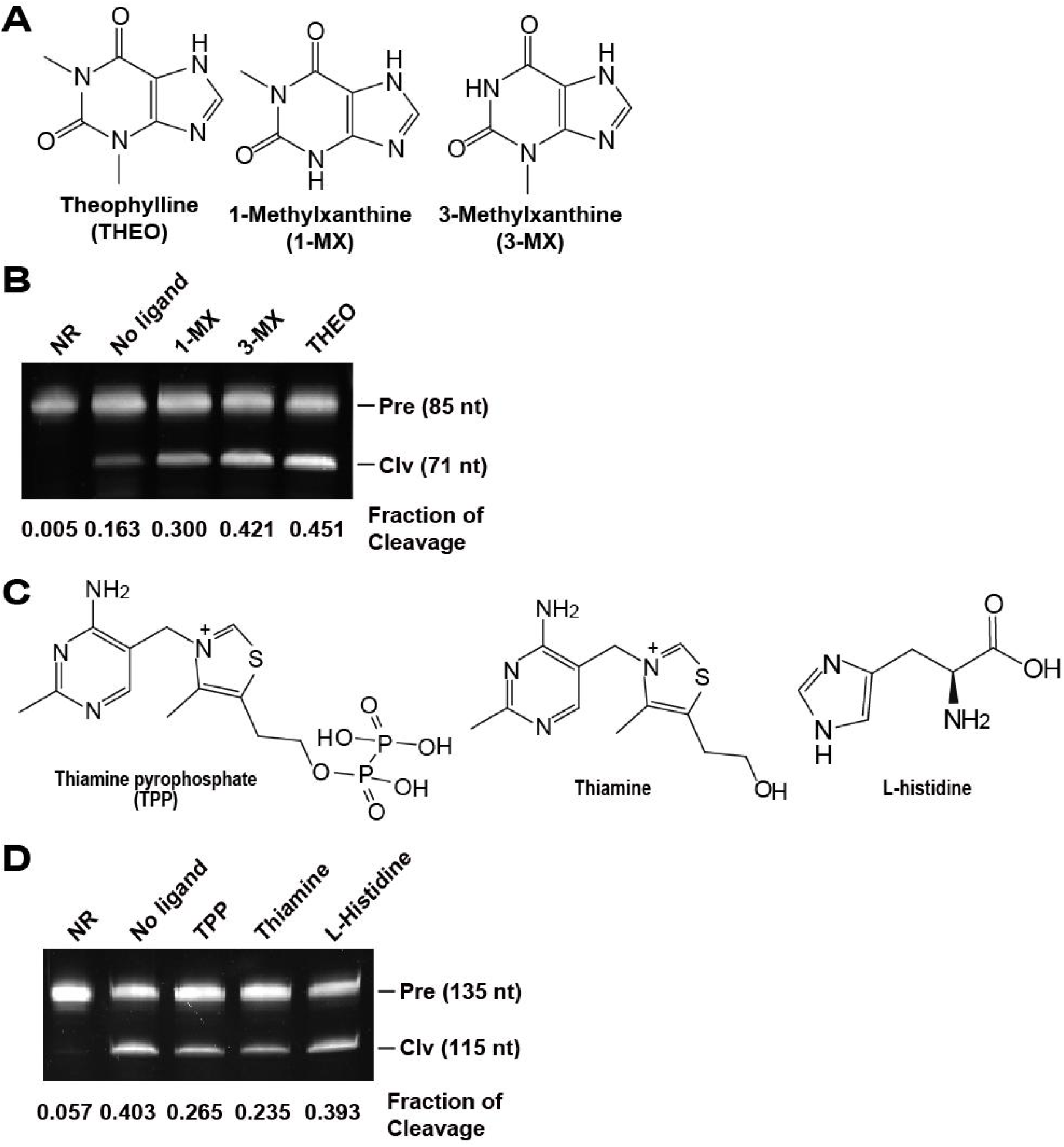
Cleavage activity of the allosteric ribozyme with different ligands. (A) Theophylline (THEO) and its analogs. (B) Induction of the cleavage of the THEO allosteric ribozyme. The concentration of the tested ligands was 100 μM in the cleavage buffer, consisting of 30 mM HEPES (pH 7.5), 100 mM KCl, and 20 mM MgCl_2_. The incubation period was set to 30 min at 25°C. (C) Thiamine pyrophosphate (TPP) and its analogs. (D) Inhibition of the cleavage of the ThiC-TPP allosteric ribozyme with a bridge of four base pairs. NR, Pre, and Clv represent no reactions (RNA only), precursors, and 5’ cleavage products, respectively. No ligand reaction contains everything necessary for ribozyme cleavage, except for ligands. The experiment was repeated thrice, and a representative gel image is shown.

### Binding affinity (*K*_D_) of the allosteric ribozymes

To investigate the binding affinity of the THEO allosteric ribozymes, we incubated the ribozyme with theophylline at concentrations ranging from 0.3 μM to 100 μM (Figure 3A). The apparent dissociation constant (*K*_D_), representing the ligand concentration required to induce or inhibit 50% of the maximum cleavage, was measured using a previously described method[42]. The *K*_D_ value for the THEO allosteric ribozyme was 3.1 ± 0.2 μM for theophylline (Figure 3B). Similar tests were conducted for other allosteric ribozymes for TPP. The *K*_D_ value for the ThiC-TPP allosteric ribozyme with a bridge of four base pairs was 1.2 ± 0.2 μM (Figure 3C and D), and the *K*_D_ value for the NCU-TPP allosteric ribozyme was 2.4 ± 1.4 μM (Figure S2). These results suggest that these allosteric ribozymes bound to their ligands tightly at the μM level and that the binding affinity could be monitored by measuring the self-cleavage products of the integrated ribozyme.

**Figure 3.**
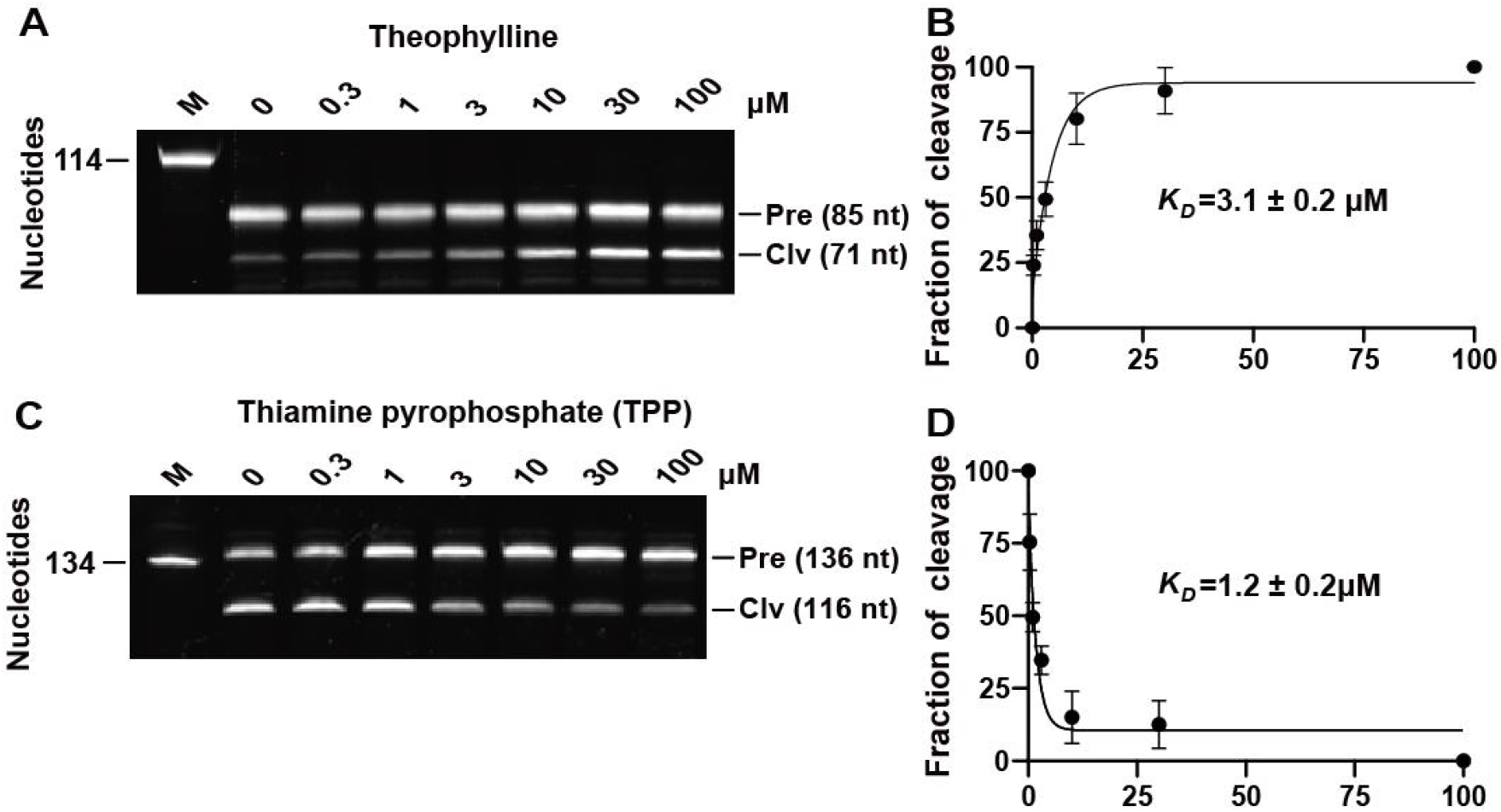
Binding affinity of allosteric ribozymes for their cognate ligands. (A) PAGE gel-based analysis of the self-cleavage of the theophylline allosteric ribozyme at different concentrations of theophylline. (B) Dissociation constant (*K*_D_) of the theophylline allosteric ribozyme. (C) PAGE gel analysis of the self-cleavage of the TPP allosteric ribozyme at different concentrations of thiamine pyrophosphate. (D) *K*_D_ of the ThiC-TPP allosteric ribozyme with a bridge comprising four base pairs. The *K*_D_ values are presented as the mean of three independent experiments with standard deviation (SD). The experiment was repeated thrice, and a representative gel image is shown. The graph was created using GraphPad Prism. If the error bar is shorter than the symbol size, Prism will not draw the bar. The other notes are the same as those listed in Figure 2.

### Effects of bridge length on allosteric ribozyme activity

The bridge length may affect the cleavage activity of the allosteric ribozymes. Although bridges with three (Figure 4) or four (Figure 1B) base pairs for the ThiC-TPP allosteric ribozyme resulted in similar cleavage activities, the *K*_D_ values varied only from 1 to 3 μM (Figures 3C and 4C). We considered that longer bridges might cause larger effects. To test this prediction, we constructed an allosteric ribozyme by integrating the hammerhead ribozyme (HHR) into the new RNA motif HMGL with bridges varying from 3 to 12 base pairs (Figure 5A). The HMGL RNA motif is a new structured noncoding RNA discovered in our bioinformatics pipeline. B**ioinformatics analyses** showed that fifty-two distinct RNA representatives of this RNA motif were found from marine sedimentary microorganisms. The structured non-coding RNA motif_HMGL was found to be located upstream of the gene encoding branched-chain amino acid-related genes. 25% of all discovered examples encode the alpha-2-isopropylmalate (IPM) synthase gene, which is the first step enzyme in the synthesis of L-leucine[60]. A further 11% of examples carry downstream genes encoding acetolactate synthase (ALS), which can catalyze pyruvate to acetolactate with high specificity and efficiency[61]. ALS is a key enzyme that catalyzes the first step in the biosynthesis of valine and isoleucine, and its activity is regulated by feedback from the products valine and isoleucine[62].We therefore speculate that this structure may sense some types of metabolites in the metabolism of branched-chain amino acids. The HMGL RNA motif could be a riboswitch candidate, so we fused an HHR ribozyme to the P1 stem for in vitro assessment. To select a proper construct for the assessment, we tested the effect of different bridge lengths on ribozyme cleavage. We constructed a construct in which two base pairs of the bridge were derived from stem II of the HHR ribozyme and the other base pairs were derived from stem P1 of the HMGL RNA motif (Figure 5A). The cleavage assay results revealed that bridges containing three or four base pairs exhibited comparable cleavage activities. In contrast, the bridge composed of six base pairs demonstrated a higher cleavage fraction, whereas the bridge composed of 12 base pairs displayed a much lower cleavage fraction (Figure 5B and C).

**Figure 4.**
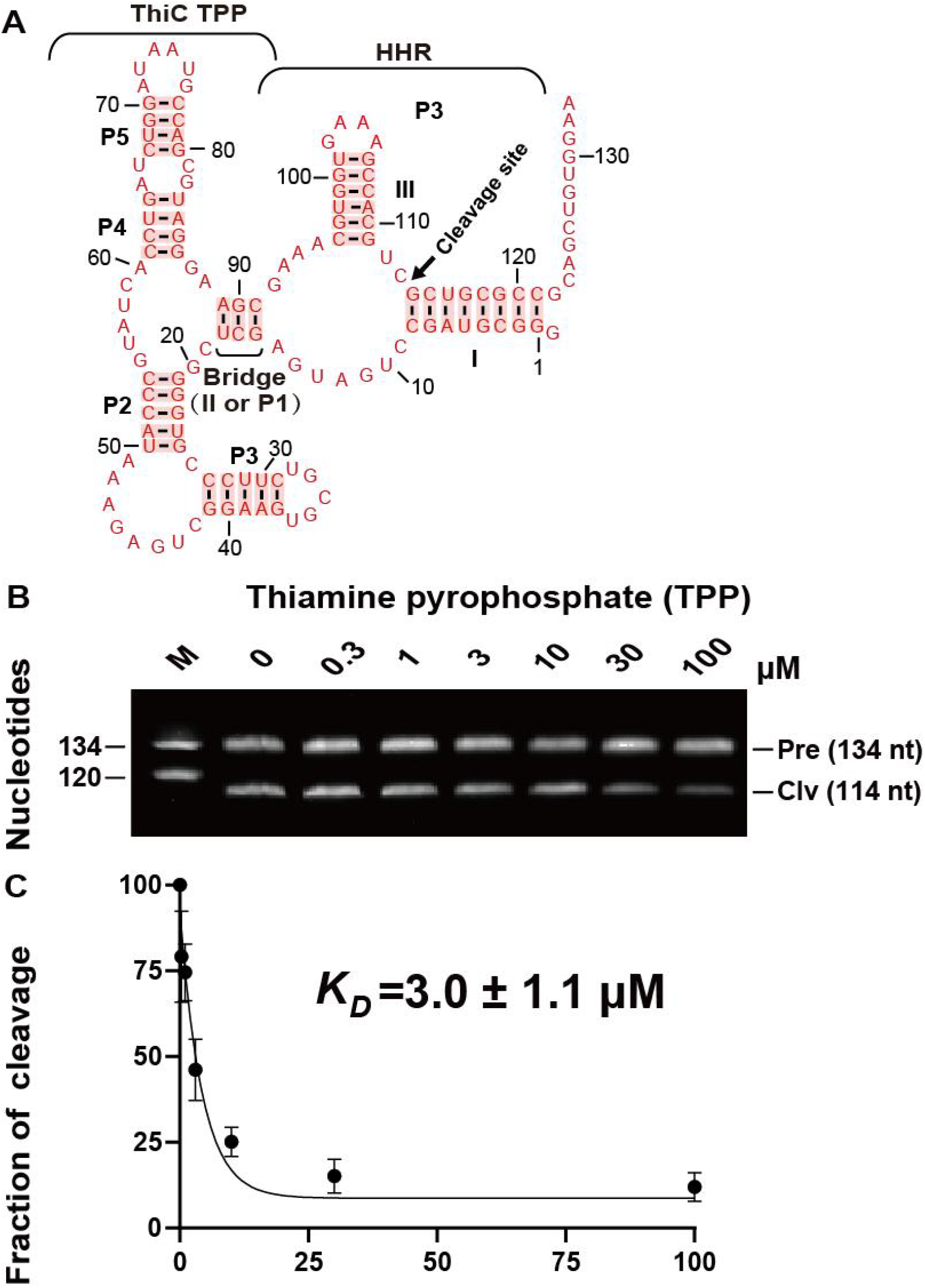
Binding affinity of the ThiC-TPP allosteric ribozymes with a bridge of three base pairs for thiamine pyrophosphate. (A) Design of the ThiC-TPP allosteric ribozyme with a bridge of three base pairs. (B) PAGE gel analysis of the self-cleavage of the ThiC-TPP allosteric ribozyme with thiamine pyrophosphate at concentrations ranging from 0 μM to 100 μM. M, Pre, and Clv represent markers, precursors, and 5′ cleavage products, respectively. (C) Dissociation constant (*K*_D_) of the ThiC-TPP allosteric ribozyme with a bridge of three base pairs for thiamine pyrophosphate.

**Figure 5.**
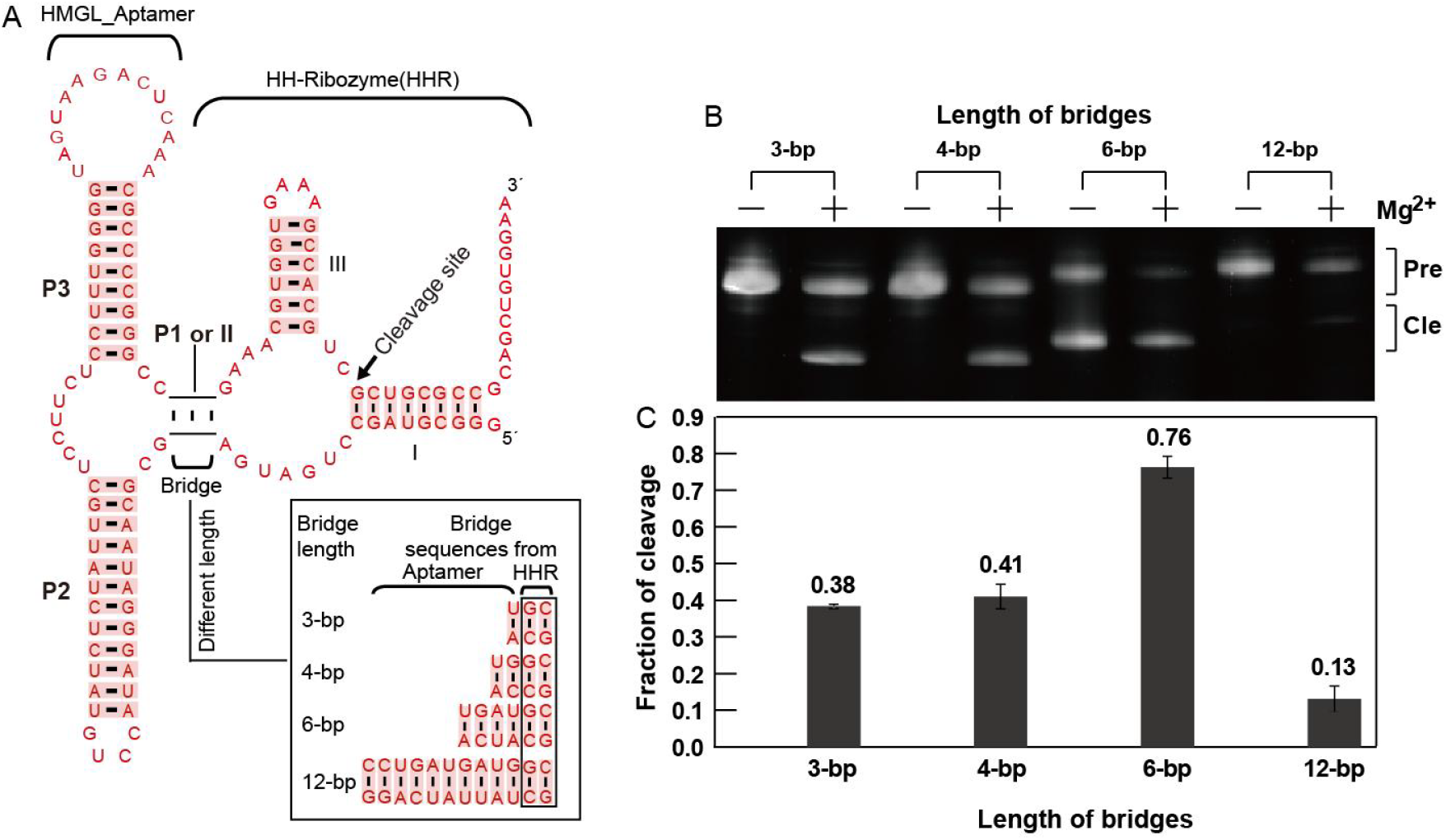
Fusion of the HMGL_RNA motif with the hammerhead ribozyme (HHR) via bridges with different base pairs of nucleotides. The bridge sequences comprise nucleotides from partial sequences of the P1 stem of the RNA motif and the II stem of the HHR.(A) Construct of the allosteric ribozyme. Two base pairs of the bridge are derived from stem II of the hammerhead ribozyme (HHR), and the other base pairs are from the P1 stem of the HMGL RNA motif. (B) Cleavage assay of the allosteric ribozyme with or without magnesium (concentration of 10 μM), where Pre represents the full length of the allosteric ribozyme, and Cle represents the 5’ cleavage product. The average fractions of cleavage for the three repeat experiments are indicated on the bar, and the standard deviations of the three repeat experiments are shown as vertical sticks on the top of each black bar.

### Function of the riboswitch candidate Motif_9307

The structured non-coding RNA Motif_9307 was found to be located in the 5’ UTR of the nitrous oxide reductase gene *nosD*[63], which encodes a subunit of nitrous oxide reductase in an operon from archaea. Sixty-five distinct RNA representatives of this RNA motif were found from archaea. The consensus model of the secondary structure of Motif_9307 (Figure 6A) shows that this motif contains P1–P5 stems in which many covariation mutations were identified by the CMfinder[55], indicating that these stems are important for the function of RNA. The enzyme reduces the greenhouse gas N_2_O to uncritical N_2_ as the final step of bacterial denitrification[63]. The NosD protein, encoded by the *nosD* gene, is a periplasmic interacting protein essential for shuttling sulfur species to the Cu_Z_ site of the 4Cu:2S cluster[56]. NosD undergoes a conformational change upon ATP hydrolysis. Consequently, the NosDFY complex alters its interaction partner, facilitating the transfer of copper ions from the chaperone NosL to the enzyme N_2_O reductase[57]. The operon was activated in response to anaerobiosis and nitrate denitrification. We predicted that the Motif_9307 RNA could sense changes in metabolite levels in cells to turn on or off the *nosD* gene.

**Figure 6.**
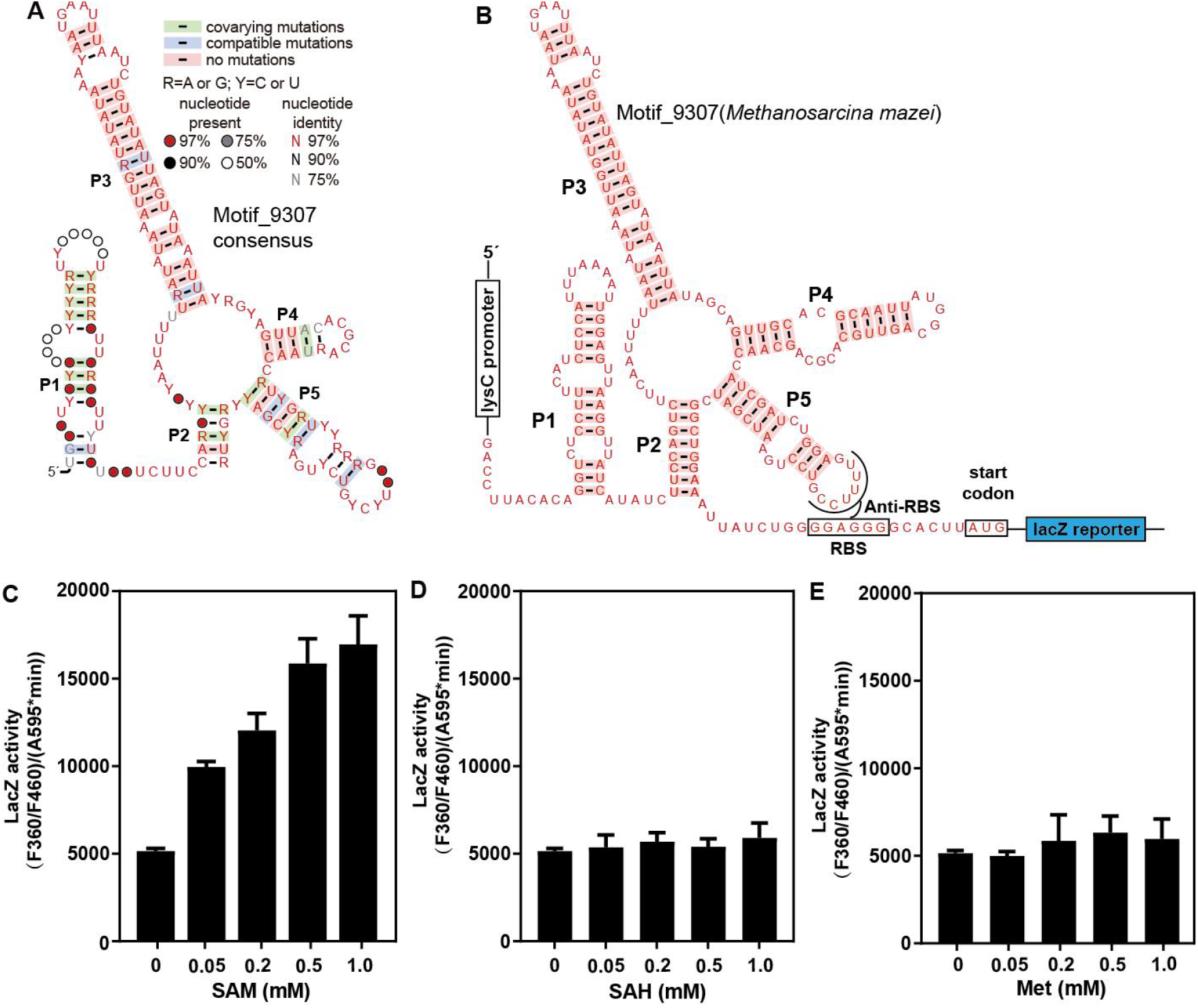
Effects of SAM, SAH, and L-methionine on *lacZ* gene expression. (A) Consensus sequence and structure of Motif_9307 based on 65 distinct RNA representatives from archaea. The structure was modeled using CMfinder to identify variations and compatible mutations in the aligned sequences and predict the secondary structure of RNA candidates. The P1–P5 stems represent base-paired substructures of RNA. Nucleotides are denoted by colored letters (R for G or A; Y for C or U) or circles (indicating the presence of any nucleotide), while colored boxes indicate evidence of natural sequence covariation or a compatible mutation that retains base pairing. (B) The construct containing the *lysC* promoter, the Motif_9307 region, and the in-frame *lacZ* gene to test the expression of Motif_9307. Effects of SAM (C), SAH (D), and L-methionine (E) on *lacZ* gene expression. The *lacZ* activities are presented as the mean of three independent experiments with standard deviation (SD). P1–P5 represent the stems of this riboswitch candidate.

To investigate whether this RNA motif can detect various metabolites, particularly certain sulfur species such as SAM, S-adenosylhomocysteine (SAH), L-methionine, and L-homocysteine, we initially constructed a plasmid that included a *lysC* promoter, the RNA motif region, and a reporter gene (*lacZ*) (Figure 6B). Subsequently, we integrated this plasmid into the genome of *B. subtilis* at the *amyE* locus. When supplemented with SAM in the range of 0.05–1.0 mM in the culture medium, **LacZ activity assay** showed that the gene expression of *lacZ* increased by approximately three-fold (Figure 6C), whereas there were no changes in gene expression when the medium was supplemented with SAH or methionine (Figure 6D and E). These results suggest that the Motif_9307 RNA can sense metabolites in cells and activate *lacZ* gene expression.

To investigate whether Motif_9307 could specifically bind to these compounds in vitro, we engineered an allosteric ribozyme by incorporating a hammerhead ribozyme into Motif_9307 RNA (Figure S3A, B, and C) through a four-base-pairing bridge (referred to as Motif_9307 allosteric ribozyme). We tested this allosteric ribozyme with various ligands, including SAM, SAH, L-methionine, L-Homocysteine, Na_2_S, glutamine, CuSO_4_, and ZnCl_2_ (Figure S3D). However, no modulations of cleavage activity were detected for these compounds (Figure S3D). In-line probing was also employed for the in vitro test, in which the ^32^P-labeled RNA was incubated with different ligands at 23°C for 36 h (Figure S4A), before separating the RNA on denatured PAGE gels. The results also showed that SAM, SAH, and L-methionine could not bind to the RNA motif directly (Figure S4B).

Subsequently, we tested the Motif_9307 allosteric ribozyme using the mixture of ligands from yeast extract (treated with chloroform to remove proteins). The results revealed that increasing the yeast extract concentration resulted in a higher cleavage fraction of the allosteric ribozyme (Figure S5A). The *K*_D_ value was determined to be 0.3 μg/μL of yeast extract (Figure S5B), suggesting that specific compounds present in the yeast extract could trigger self-cleavage of the allosteric ribozyme.

## Discussion

### Integration of the ribozyme into aptamers through a bridge to measure ligand-binding events

Several approaches can be applied to measure ligand binding. In-line probing can not only detect the *K*_D_ value but can also deduce which nucleotides are important for ligand binding[44]. ITC can be used to detect the binding of structured RNA to the ligand by measuring the temperature shift during the binding process[46]. SYBR Green can be used to analyze binding by detecting the competition between SYBR Green molecules and the ligand[47]. However, these methods involve a long and difficult procedure to crystallize RNA, use radiative elements, prepare a large amount of pure RNA, or deal with insensitive measurements. Here, we designed a bridge to integrate the ribozyme into an aptamer (or riboswitch candidates) to monitor the ligand-binding events by measuring the cleavage of the allosteric ribozyme. We found this method to be very sensitive for measuring ligand-binding events by detecting ribozyme cleavage activity.

This approach only requires approximately 1 pmol of allosteric ribozyme RNA and from nM to μM of ligands, which are the same amounts used for in-line probing. Using our approach, the measured *K*_D_ values for the THEO, ThiC-TPP, and NCU-TPP allosteric ribozymes were 3.1 μM, 1.2 μM, and 2.4 μM, respectively (Figures 3 and S2). Similarly, the *K*_D_ value for the theophylline aptamer measured using equilibrium filtration analysis with ^14^C-labeled theophylline was 3.1 μM[58], and the *K*_D_ value for the ThiC-TPP riboswitch with a long construct from bacteria was 0.1 μM by in-line probing with ^32^P-labeled RNA[59]. These results indicate that the *K*_D_ values measured using our approach are close to those measured using in-line probing and equilibrium filtration analysis. Moreover, as our method does not require radioactive elements to label RNAs or ligands, it is safer and more convenient.

### Various ribozymes can be used for integration with an appropriate bridge

The hammerhead ribozyme used in this study has two versions; one has short stem I and stem III (Figure 1A), while the other has slightly longer stem I and III (Figure 1B). However, both versions worked well in the experiments. Whether these versions of hammerhead ribozymes have different effects on the same aptamers or riboswitches has not been tested. Recently, modular aptazymes with the full-length hammerhead ribozyme have been reported, wherein similar but short communication modules are used[64]. We believe that other ribozymes, such as the pistol and twister ribozymes, can also be integrated as long as they can be properly merged with aptamers. For DNA aptamers, DNAzymes such as I-R1 or II-R1 DNAzymes [65] can be used to measure the ligand binding for DNA aptamers after integration into the aptamers[42, 43].

### Function and design of bridges in allosteric ribozymes

The bridge can transfer the ligand-binding signals from the aptamer to the ribozyme. The ligand binding causes the conformational change of the aptamer and results in the twisting or folding of the bridge at the same time. If the twisting or folding of the bridge is in favor of proper ribozyme folding, ribozyme cleavage is induced; otherwise, it is inhibited. These predictions are exactly what we observed for the theophylline and TPP allosteric ribozymes (Figure 3). The cleavage of the theophylline allosteric ribozyme was induced by theophylline binding, whereas the cleavage of the TPP allosteric ribozyme was inhibited by TPP binding.

To keep both the aptamer and the ribozyme intact during integration, we designed the bridge to contain several base pairings from stem P1 of the aptamer and stem II of the ribozyme (Figure 1, Figure 4 and Figure 5). The composition of the bridge needs to be adjusted to ensure that the ribozyme can self-cleave to some degree. For optimization of bridge length, we used bridges of three to four base pairs (or even longer) for each allosteric ribozyme and found that some constructs had better performance (better cleavage rates and ligand reactions), and we reported the optimal constructs for each allosteric ribozyme (Figure 1, Figure 5 and Figure S1). In other early research, the bridges of four or even one base pair could function well for the ligand sensing of the allosteric ribozyme composed of the theophylline aptamer and the hammerhead ribozyme[66, 67]. In our experiments, we found that a bridge of three to six base pairs usually could maintain good cleavage of the allosteric ribozymes, whereas too long bridges may lower the cleavage activity of the ribozymes (Figure 5).

In addition, when testing the bridge length, we found that the HMGL allosteric ribozyme with a six-base pair bridge showed self-cleavage even in the absence of Mg^2+^ (Figure 5). We do not know the reason for this. It has been shown that hammerhead, hairpin and VS ribozymes do not require Mg^2+^ for their enzyme activity, NH_4_^+^ and monovalent metal ions can also support their enzyme activity. We thought that this was most likely due to the unique structure of the allosteric ribozyme with a six-base pair bridge and the presence of 100 mM KCl in our cleavage buffer causing the cleavage. This requires further investigation.

### Application of this approach to measuring binding ligands of riboswitch candidates and other RNA aptamers

Since the invention of the in-line probing method by Dr. Breaker’s group, it has been used to search for ligands for riboswitch candidates. For example, it has been used for the discovery of a fluoride riboswitch that can sense toxic fluoride in the cell[68]. Many other riboswitch candidates, such as the fluoride riboswitch, c-di-GMP riboswitch, guanine riboswitch, and TPP riboswitch, have been confirmed using this method in Dr. Breaker’s laboratory[1, 2, 69, 70]. However, many groups do not conveniently use in-line probing because it requires the use of the radioactive element ^32^P. ITC requires a large amount of RNA and expensive equipment for the measurement, while the SYBR Green fluorometric assay was not sensitive under the tested conditions. Therefore, we believe that this ribozyme integration method can be used as an alternative approach for monitoring ligand-binding events.

In addition, Dr. Breaker’s lab has constructed several allosteric ribozymes by selecting communication modules (CMs) from random sequences using SELEX to make allosteric ribozymes and has shown that ribozyme activity can be regulated by a riboswitch or an aptamer[38, 39]. Our main goal is to construct allosteric ribozymes to measure ligand binding events by designing linker bridges to connect riboswitches or aptamers to a ribozyme. The bridges are usually composed of one to two base pairs from stem II of the hammerhead ribozyme and two to three base pairs from one of the stems of a riboswitch or an aptamer (Figure 1, Figure 4 and Figure 5). This design requires the testing of several bridges to select the one with adequate ribozyme cleavage activity for monitoring ligand binding events, rather than using SELEX to obtain communication modules.For the 9307_motif allosteric ribozyme test, although we observed some modulations in the experiment. Further experiments are needed, such as chromatographic column separation of yeast extract and testing each subset separately, to definitively identify which component is responsible for the modulation.

In summary, we designed a bridge to join a hammerhead ribozyme with riboswitch candidates or aptamers to observe and measure ligand-binding events based on the induction or inhibition of ribozyme cleavage. After testing the approach with the theophylline aptamer, two TPP riboswitches, and one riboswitch candidate, we found that the approach represents an alternative method for measuring ligand-binding events. Combined with in vivo data, our findings suggest that the riboswitch candidate Motif_9307 functions as a metabolite-sensing riboswitch, potentially regulating gene expression of *nosD*. Furthermore, one or more compounds in the yeast extract may serve as ligands for this riboswitch candidate.

## Acknowledgements

We thank LetPub (www.letpub.com.cn) for its linguistic assistance during the preparation of this manuscript.

## Data availability

The data underlying this article are available in the article and in its online supplementary material.

