## Supplementary material for "Integration of the hammerhead ribozyme into structured RNAs to measure ligand-binding events for riboswitch candidates and aptamers": Integration of the hammerhead ribozyme into structured RNAs_suppl_2025_3-30-revised.pdf

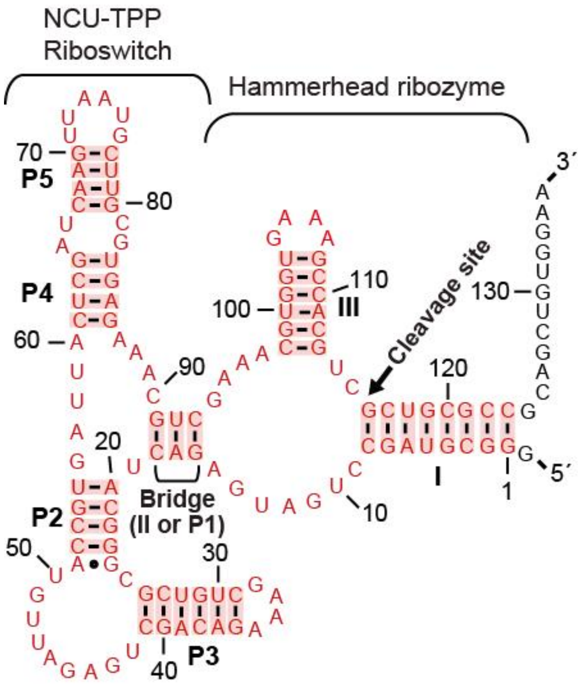


Figure S1 Construction of the NCU-TPP allosteric ribozyme with a bridge of three base pairs. We fused the TPP riboswitch, derived from the fungal gene *NCU01972*, with the hammerhead ribozyme to create a novel construct called the NCU-TPP allosteric ribozyme. This chimeric structure consisted of a bridge comprising three base pairs, which replaced the P2 stem of the hammerhead ribozyme. The stems of the hammerhead ribozyme were denoted as stems I, II and III, while those of the riboswitch were labeled as P1, P2, P3, P4, and P5.

删除[Windows User]: T

删除[Windows User]:

删除[Windows User]: was integrated

删除[Windows User]: , resulting in the

删除[Windows User]: creation

删除[Windows User]: of

删除[Windows User]: known as

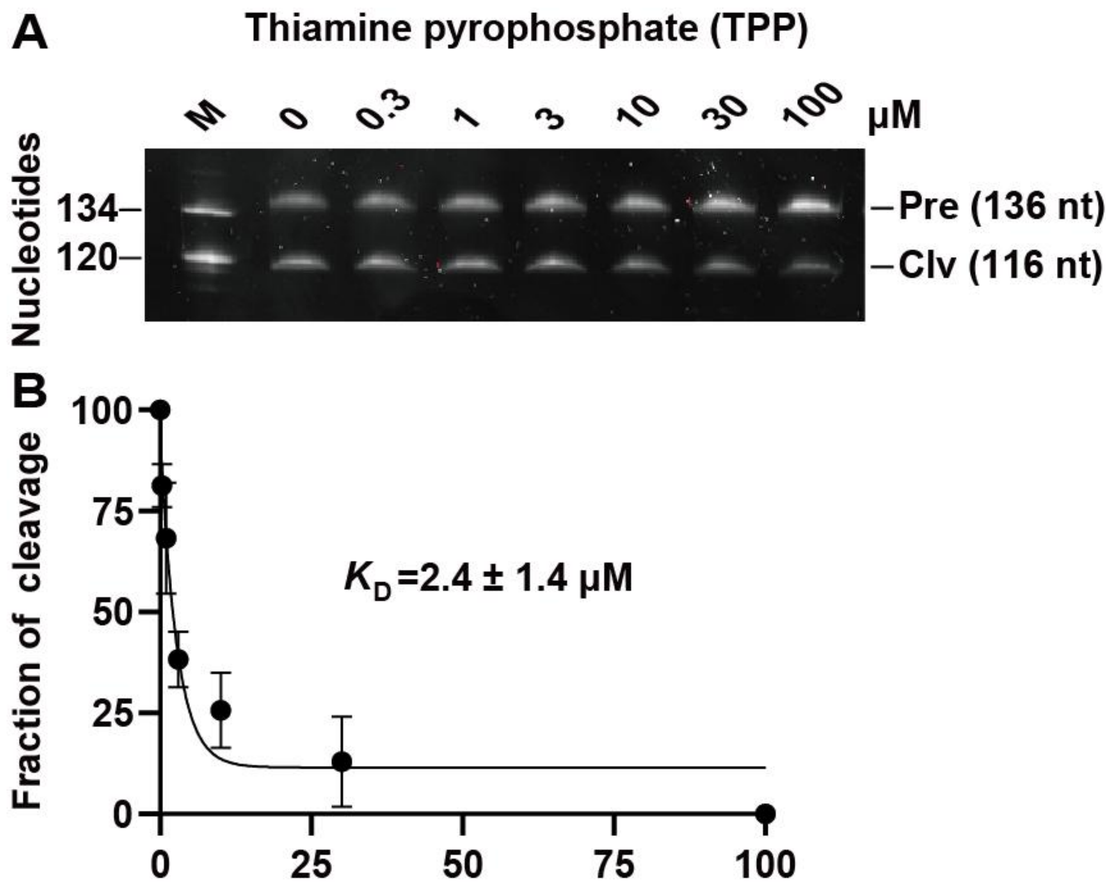


Figure S2 Binding affinity of the NCU-TPP allosteric ribozymes with a three-base pair bridge for thiamine pyrophosphate. (A) PAGE gel analysis of the self-cleavage of the NCU-TPP allosteric ribozyme with thiamine pyrophosphate at concentrations ranging from 0.3 μM to 1000 μM. M, Pre, and Clv represent markers, precursors, and 5' cleavage products, respectively. (B) Dissociation constant ($K_D$) of the NCU-TPP allosteric ribozyme for thiamine pyrophosphate. The $K_D$ values are the mean of three independent experiments with standard deviation (SD).

删除[Windows User]: of three base pairs

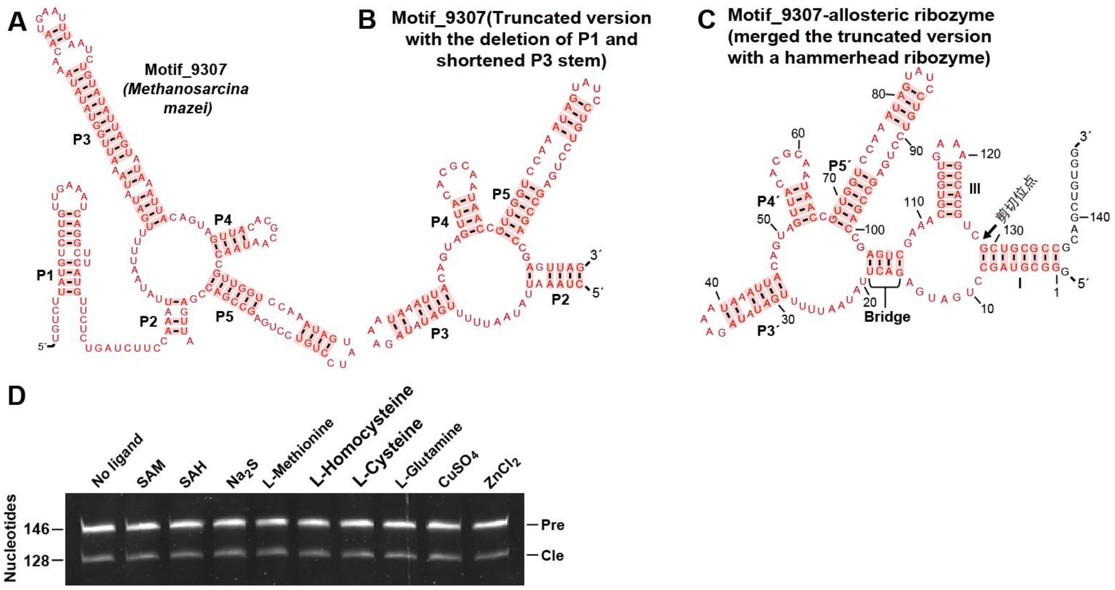


Figure S3 Effect of different ligands on the cleavage of the 9307_motif allosteric ribozyme. The concentration of the ligands SAM (S-adenosine-methionine), S-adenosine-homocysteine (SAH), L-methionine, L-homocysteine, glutamine, $CuSO_4$, $ZnCl_2$, and $Na_2S$ was 1 mM. (A) The secondary structure of one of the Motif_9307 representatives from *Methanosarcina mazei*. (B) To fuse Motif_9307 with the ribozyme, we deleted the P1 stem P1, truncated the P3 stem of the Motif_9307 RNA and renamed the stems from p3' to P5'. (C) The fusion of Motif_9307 with the ribozyme through a four-base pairing bridge, consisting of three base pairs from the aptamer and one base pair from the hammerhead ribozyme. (D) Cleavage of the allosteric ribozyme in the presence of different ligands. No ligand represents the reaction without the addition of any ligands. Other notes are the same as those listed in Figure S2.

删除[Windows User]: and

删除[Windows User]: of four base pairings

删除[Windows User]: comprising

删除[Windows User]: The c

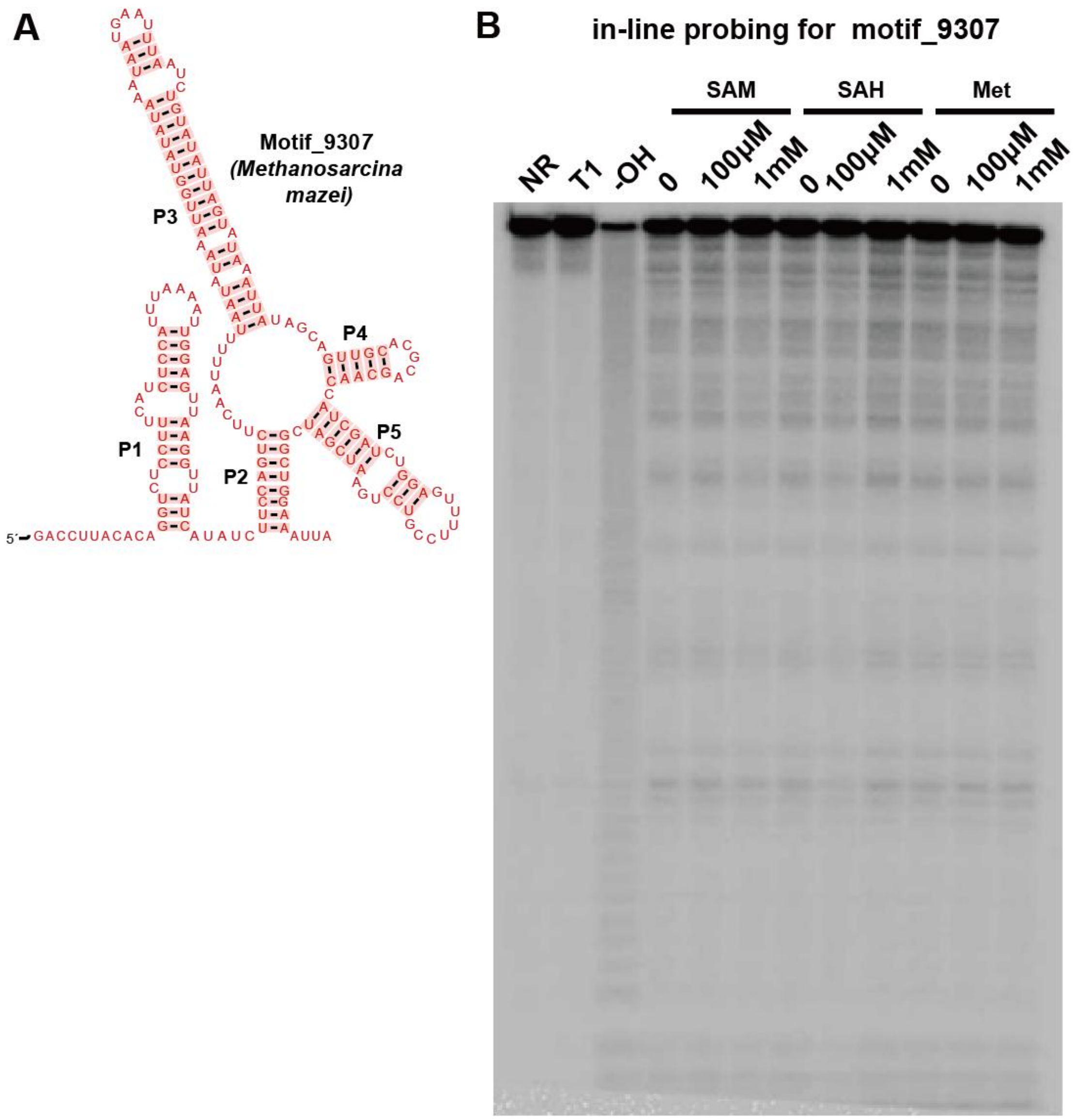


Figure S4 In-line probing assay for Motif_9307. (A) Sequence and secondary structure of a representative from Motif_9307 in *Methanosarcina mazei*. (B) In-line probing assay for the representative RNA of Motif_9307 from *Methanosarcina mazei.* PAGE gel analysis shows in-line probing reactions of the 5' $^{32}P$-labeled Motif_9307 RNA with SAM, SAH, or L-methionine (Met) at concentrations ranging from 0 to 1000 μM. NR, T1, and $^{-}OH$ designate no reaction, partial digestion with RNase T1, and partial digestion with alkali, respectively.

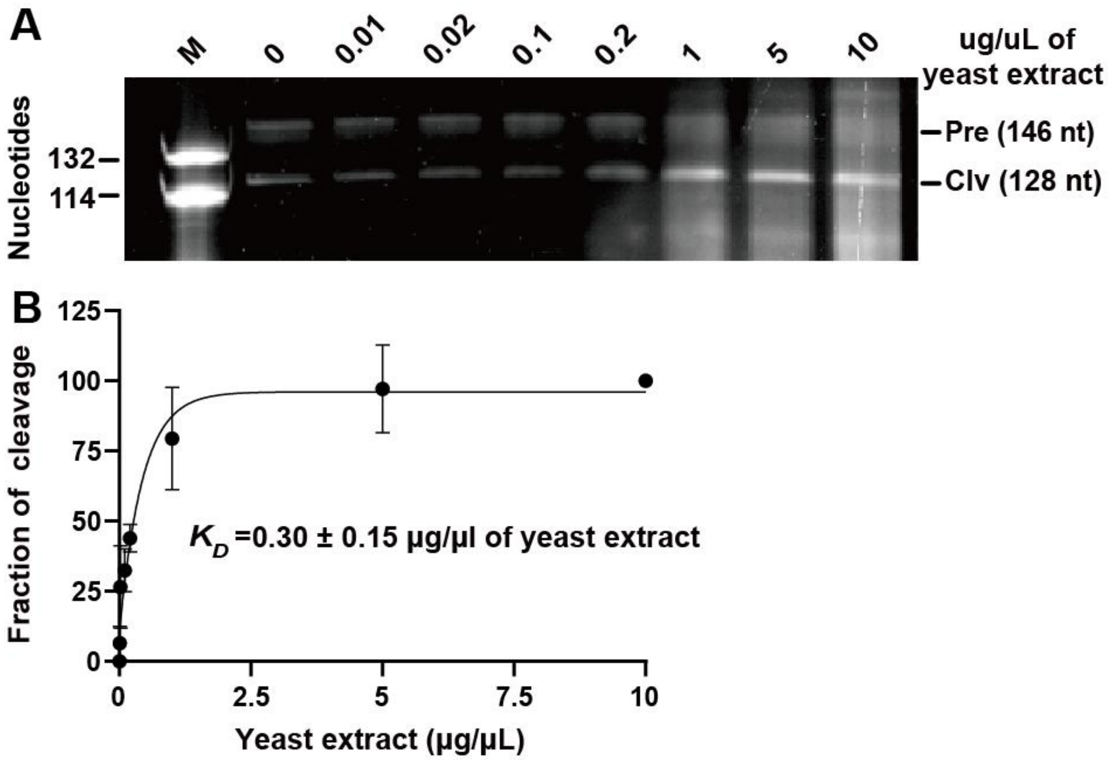


Figure S5 Induction of self-cleavage of the Motif_9307 allosteric ribozyme by yeast extract. (A) PAGE gel analysis of the Motif_9307 allosteric ribozyme with different concentrations of yeast extract. The experiment was repeated three times, and one representative gel image is shown. (B) $K_D$ of the Motif_9307 allosteric ribozyme for the yeast extract. The error bars represent the standard deviation for each concentration of the yeast extract. If the error bar is shorter than the symbol size, Prism will not draw the bar. Other notes are the same as those listed in Figure S2.

Table S1 Primers and oligos

| Name | Sequences | Notes |
|---|---|---|
| Marker M1 (132 nt) | 5'-TAATACGACTCACTATAGGGAAACTGTCCTTCCCGGAAGGTTTCAAATGGGAACGTGTTATGAACTTCGAAGACGGTGGTGTTGTTACCGTTACCCAGGACTCCTCCCTGCAAGACGGTGAGTTCATCTACAAAGTTAAACTGCGTGGTAC | The DNA template contains a T7 promoter to make the 132 nt RNA marker (the fragment is part of the gene for red fluorescent protein). |
| Marker M2 (120 nt) | 5'-TAATACGACTCACTATAGGGAAACTGTCCTTCCCGGAAGGTTTCAAATGGGAACGTGTTATGAACTTCGAAGACGGTGGTGTTGTTACCGTTACCCAGGACTCCTCCCTGCAAGACGGTGAGTTCATCTACAAAGTTAA | The DNA template contains a T7 promoter to make the 120 nt RNA marker (the fragment is part of the gene for red fluorescent protein). |
| Marker M3 (114 nt) | 5'-TAATACGACTCACTATAGGGAAACTGTCCTTCCCGGAAGGTTTCAAATGGGAACGTGTTATGAACTTCGAAGACGGTGGTGTTGTTACCGTTACCCAGGACTCCTCCCTGCAAGACGGTGAGTTCATCTACAA | The DNA template contains a T7 promoter to make the 114 nt RNA marker (the fragment is part of the gene for red fluorescent protein). |
| Marker M4 (87 nt) | 5'-GAAACTGTCCTTCCCGGAAGGTTTCAAATGGGAACGTGTTATGAACTTCGAAGACGGTGGTGTTGTTACCGTTACCCAGGACTCCTC | The DNA template contains a T7 promoter to make the 87 nt RNA marker (the fragment is part of the gene for red fluorescent protein). |
| Theophylline I-forward | 5'-TAATACGACTCACTATAGGGCGTAGCCTGATGAGCCCTAATACCAGCCG | Use theophylline I-forward and I-reverse for |

| | | |
|---|---|---|
| | AAAGGCCCTTGGCAG | overlapping PCR to make a DNA template for the theophylline allosteric ribozyme. |
| Theophylline I-reverse | 3'-CCACAGCTCGCAGCGACGTGTTTCCACGTTTCGCTCTACTGCCAAGGGCCTTTCG | |
| Theophylline II-forward | 5'-TAATACGACTCACTATAGGGCGTAG | Use II-forward and II-reverse short primers to amplify the full-length DNA. |
| Theophylline II-reverse | 3'-CCACAGCTCGCAGCGA | |
| ThiC-TPP I-forward | 5'-TAATACGACTCACTATAGGGCGTAGCCTGATGAGACTCGGGGTGCCCTTCTGCGTGAAGGCTGAGAAATACCCGTATCACCTGATCTGG | Use ThiC-TPP I-forward and I-reverse for overlapping PCR to make a DNA template for the ThiC-TPP allosteric ribozyme. |
| ThiC-TPP I-reverse | 3'-TTCCACAGCTGCGGCGCAGCGACGTGGCTTTCACCACGTTTCGACTTCCCTACGCTGGCATTATCCAGATCAGGTGATACGGGTATTTC | |
| ThiC-TPP-II-forward | 5'-TAATACGACTCACTATAGGGCGTAG | Use II-forward and II-reverse short primers to amplify the full-length DNA. |
| ThiC-TPP-II-reverse | 3'-TTCCACAGCTGCGGCGCAG | |
| NCU-TPP-I-forward | 5'-TAATACGACTCACTATAGGGCGTAGCCTGATGAGACTACGGGCGCTGTCGAAAGACAGCTGAGATTGTACCGTGATTACTCGATCAAG | Use NCU-TPP I-forward and I-reverse for overlapping PCR to make a DNA template for the NCU-TPP allosteric ribozyme. |
| NCU-TPP-I-reverse | 3'-TTCCACAGCTGCGGCGCAGCGACGTGGCTTTCACCACGTTTCGACGTTTCTCACGCAAGCATTAACTTGATCGAGTAATCACGGTACAATC | |

| | | |
|---|---|---|
| NCU-TPP-II-forward | 5'-TAATACGACTCACTATAGGGCGTAG | Use II-forward and II-reverse short primers to amplify the full-length DNA. |
| NCU-TPP-II-reverse | 3'-TTCCACAGCTGCGGCGCAG | |
| Motif_9307-I-forward | 5'-GGCGTAGCCTGATGAGACTTATAATTTTTGATATAGAAATAAATTACAGTAGTTACACGCAATAACCGTTGGTCCAAATAGTATCCTG | Use I-forward and I-reverse for overlapping PCR to make a DNA template for the motif_9307 allosteric ribozyme. |
| Motif_9307-I-reverse | 3'-TTCCACAGCTGCGGCGCAGCGACGTGGCTTTCACCACGTTTCGACTCGGTCGGCTCAGGACAGGATACTATTTGGACCAACGG | |
| Motif_9307-II-forward | 5'-TAATACGACTCACTATAGGGCGTAGCCTGATGAGACT | Use II-forward and II-reverse to amplify the full-length Motif_9307 allosteric ribozyme in which the T7 promoter was added to the 5' end. |
| Motif_9307-II-reverse | 3'-TTCCACAGCTGCGGC | |
| Partial sequence of pBS1ClacZ (reconstructed region between EcoRI and SalI) | 5'-GAATTCTGCAAAAATAATGTTGTCCTTTTAAATAAGATCTGATAAAATGTGAACTAATGGATCCACAGTACATAAAAAAGGAGACAAGCTTACGATGGTCGTTTTACAACGTGACTGGGTCGAC | The DNA template contains EcoRI, BamHI, HindIII, and SalI for insertion of RNA motifs that are in-frame with the lacZ reporter. |
| Lysc_F | 5'-GATCTGATAAAATGTGAACTAATGGATCC | Use Lysc_F and Lacz_R for PCR. True transformants should contain this fragment. |
| Lacz_R | 5'-GCAGCAACGAGACGTCAC | |
| Amp_F | 5'- GACTTGGTTGAGTACTCACCAG | Use Amp_F and Ori-R for |

| | | |
|---|---|---|
| Ori-R | 5'- GCAGAGCGAGGTATGTAGG | PCR. True transformants should not contain this fragment. |
